# Native Korean wild mice (*Mus musculus*): molecular phylogeny and morphometrics

**DOI:** 10.1101/2024.05.06.592665

**Authors:** Daewoo Kim, Jooseong Oh, Jang Geun Oh, Hee-Young Yang, Geun-Joong Kim, Tae-Hoon Lee, Bae-Geun Lee, Chungoo Park, Dong-Ha Nam

**Affiliations:** Department of Biological Sciences, College of Natural Sciences, Chonnam National University, Gwangju, Korea; School of Biological Sciences and Technology, College of Natural Sciences, Chonnam National University, Gwangju, Korea; Jeju Wildlife Problem Institute, Jeju-do, Korea; Preclinical Research Center, Daegu-Gyeongbuk Medical Innovation Foundation (KMEDIhub), Daegu, Korea; Department of Oral Biochemistry, Dental Science Research Institute, Korea Mouse Phenotype Center (KMPC), School of Dentistry, Chonnam National University, Gwangju, Korea; National Institute of Ecology, Seocheon-gun, Chungcheongnam-do, Korea

**Keywords:** House mouse, taxonomy, phylogeny, morphology, Korean Peninsular

## Abstract

Taxonomic status of house mice in the Korean Peninsula remains poorly understood. Here, we analyze genetic and morphological characteristics of mice from Korea and evaluate their phylogenetic relationships to the well-known primary subspecies. Using a comprehensive set of publicly available genetic data (mtDNA *cytb* gene), Korean mice including our specimens from islands, mountains, and agricultural fields were identified to *Mus mus musculus*. External morphology, such as tail ratios of our specimens, resembled previously assigned subspecies (e.g., *M. m. molossinus*, *M. m. utsuryonis*, and *M. m. yamashinai*), suggesting a single subspecific group within *M. m. musculus*. Korean mice displayed a distinctive landmark configuration around the snout, with a relatively short and slender premaxillary tooth-patch width (PMXW) and a larger maxillary tooth-row length (MXTL) compared to laboratory strains derived from *M. m. domesticus*. Our investigation provides insights into the phylogenetic relationships and taxonomic status of Korean mice relative to the primary lineages of *M. musculus* subspecies. Understanding the evolutionary history of Korean *M. m. musculus* sheds new light on how their spatiotemporal dynamics led to diversification, with the Korean Peninsula serving as an ecological bridge between East Eurasia and neighboring regions.

## Introduction

Wild house mice (*Mus musculus*) are human commensals that have expanded their ecological niches alongside human activities, facilitating the continuous accumulation of morphological and genetic variations. Emerging from the Indian subcontinent, they subsequently dispersed globally and diverged into three primary subspecies: *M. m. castaneus* (CAS) present across Southeast Asia, *M. m. domesticus* (DOM) ranging from Western Europe through Africa to South and North America, and *M. m. musculus* (MUS) that has expanded its ranges throughout Eurasia [1].

The Korean Peninsula, located in the Far East of Eurasia, presents ecological barriers that influence mammal distribution and population dynamics. Previous studies using external body measurements and fur/hair color phenotypes classified wild mice in the Korean Peninsula into three subspecies: *M. m. molossinus* [2,3], *M. m. utsuryonis* [4], and *M. m. yamashinai* [5]. However, this classification based on subtle morphological variations remains controversial, as it may not accurately reflect the true evolutionary relationships (phylogeny) of these mice. Given that MUS in East Eurasia might have entered the Korean Peninsula from northern China, with human immigration presumably facilitating its geographical dispersal [6,7], the question remains whether the assigned subspecies of mice truly reflect their taxonomic positions [8–14].

We integrate existing genetic and morphological data from house mice with mtDNA makers, 3D micro-computed tomography (CT)-scanned landmark and craniometrics to address the evolutionary relationships of house mice in the Korean Peninsula. This approach clarifies the phylogenetic relationships of these mice to the three primary house mouse subspecies.

## Materials and Methods

### Samples

From 2017 to 2023, we collected wild mice from various locations across Korea, including islands (Gageo-do, Jeju-do, and Ulleung-do; n = 6), mountains (Yeongju-si and Wanju-gun; n = 3), and agricultural fields (Goesan-gun, Goheung-gun, and Naju-si; n = 14) (Fig 1). In addition to the wild mice, we used four commercially available laboratory mouse strains (CBA, C57BL/6H, C3H, and BALB/c derived from DOM) and our inbred lines from Goesan-gun (KG01BRA/1CNUf1∼f3 and KG01BRA/1CNUm1∼m3) (Each group included three male and three female mice, aged 9 weeks.). The animal study was approved in accordance with the regulations by the Chonnam National University Institutional Animal Care and Use Committee (CNU IACUC-YB-2021-103).

**Fig 1.**
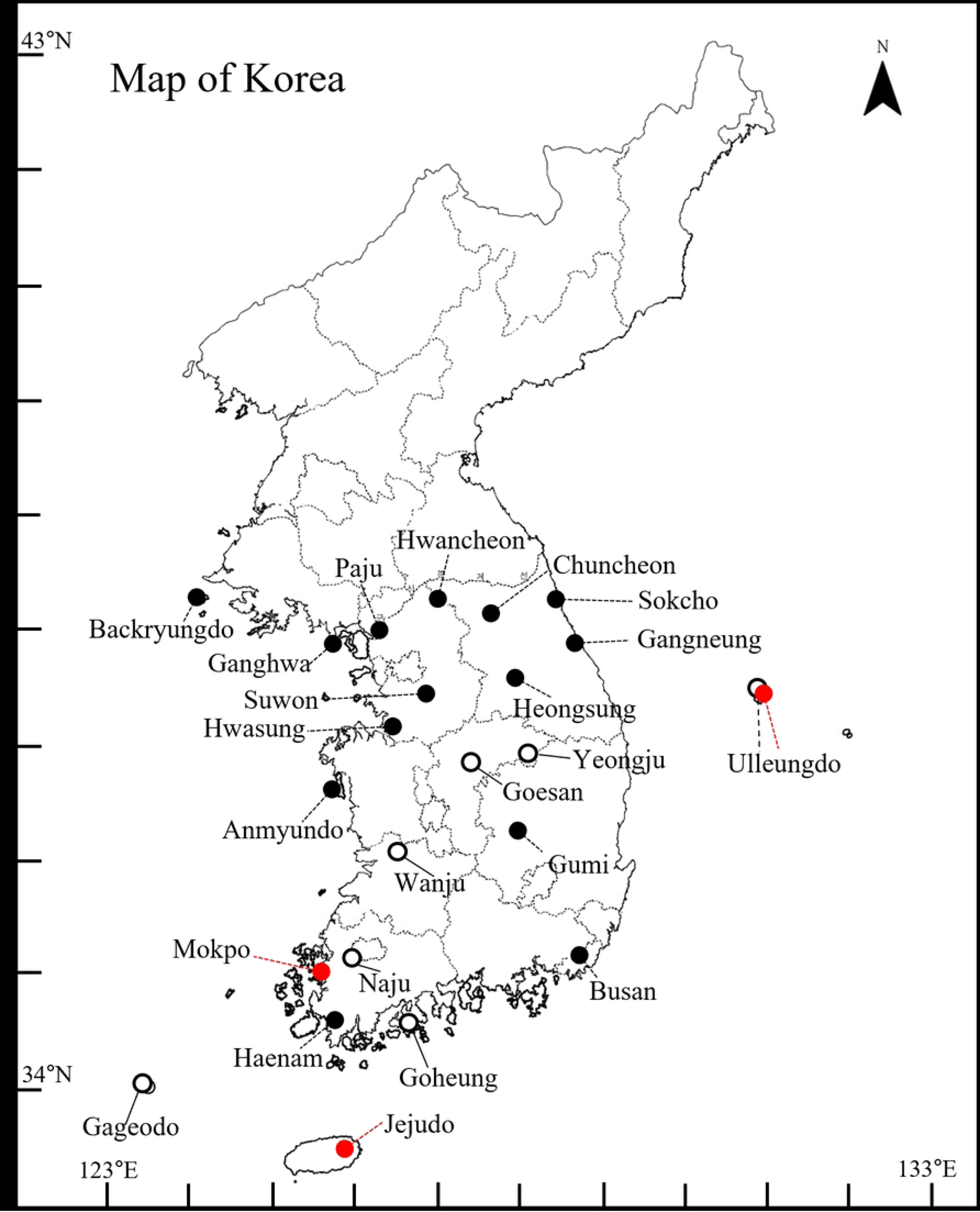
Sampling locations of Korean house mice in the present study (open circle). Red circles, three subspecies have been assigned morphologically as *M. m. molossinus* on Jeju island, *M. m. utsuryonis* on Ulleung island, and *M. m. yamashinai* on the Korean Peninsula including Mokpo [2–5]; solid black circles, previously studied locations for Korean mice lineage (Table 4).

### Molecular analysis

Genomic DNA was extracted from approximately 0.5 cm of tail tip, obtained by scraping the skin with a sterilized scalpel to remove surface contaminants, from 23 specimens. The mitochondrial cytochrome b gene (*cytb*) (1,140 bp) was amplified by PCR using universal primers (L7, ACC AAT GAC ATG AAA AAT CAT CGT T; H6, TCT CCA TTT CTG GTT TAC AAG AC). PCR amplification conditions for *cytb* were as follows: initial denaturation at 95 °C for 5 minutes, followed by 36 cycles of denaturation (95 °C for 40 seconds), annealing (56 °C for 60 seconds), and extension (72 °C for 60 seconds). A final extension step was performed at 72 °C for 5 minutes. The purified PCR products were sequenced using the BigDye v3.1 (ABI).

We retrieved publicly available *cytb* sequences for *M. musculus* subspecies, along with representative sequences of a closely related species (e.g., *M. spicilegus*) as an outgroup. Our mtDNA sequence data for each region were deposited in GenBank with accession numbers OQ506519.1 ∼ OQ506540.1 (S1 Table). We used MEGA11 to align sequence data from mouse subspecies in wild populations and wild-derived strains. The number of mouse populations were used for AMOVA (analysis of molecular variance) and calculation of Nei’s genetic distance based on *p*-distance using the MEGA11 program. From the AMOVA, we obtained the fixation index (*Fst*), which describes the processes leading to genetic differentiation among and within populations. Haplotype networks for 93 *cytb* sequences were constructed using the median-joining network (S1 Table) to illustrate potential clustering of the main network topology as a reticulation split. Maximum Likelihood (ML) and Bayesian Inference (BI) phylogenies were constructed for each *cytb* data set (S1 Table) with the optimized Hasegawa-Kishino-Yano substitution model. ML bootstrap (BS) analysis and BI posterior probabilities (PP) were performed with 1,000 replications using MEGA11. A sub-group was clustered within each of the major mtDNA lineages only if there was bootstrap support of >0.7 from the ML analysis, combined with concordant structure in the haplotype networks. To assess demographical history, the number of haplotypes (*H*), haplotype diversity (*Hd*), and nucleotide diversity (*π*) were evaluated using DnaSP software. Tajima’s *D* and Fu’s *Fs* values were also estimated with ARLEQUIN software.

### Morphometrics

The micro-CT scanning system captures elaborate tomographic images of specimens. The specific procedures were described previously [15,16] with some minor adjustments. For micro-CT scanning, each mouse was placed on an acrylic plate attached to an animal bed inside the micro-CT scanner. To ensure accurate bone length measurements in the resulting images, the vertebral column was carefully stretched, and the head, body, and tails were gently pressed onto the plate. The bone structures of the specimens were examined using a micro-CT scanning system combined with a Quantum GX micro-CT imaging system (PerkinElmer). A 3D micro-CT scanning system equipped with an X-ray tube (voltage 90 kV, current 80 mA) was used to scan the mouse skeleton. For whole-body scans, a voxel size of 90 μm and a field of view (FOV) of 45 mm were used, with a working distance of 108 mm and a scan time of 2 minutes in high-resolution scan mode. Cranial images were acquired using a voxel size of 50 μm, FOV of 25 mm, working distance of 55 mm, and scan time of 2 minutes in standard scan mode. The scanned skeletal data were reconstructed into 3D tomograms comprising high-contrast images of the skeletal parts of interest.

To quantify both size and shape of the geometric objects, we used Micro-CT Viewer (Quantum GX, PerkinElmer) and ImageJ software. We measured the lengths of the following skeletal parts from wild mice (n = 13), 4 laboratory mouse strains (CBA, C57BL/6H, C3H, and BALB/c derived from DOM) (n = 6 each), and our inbred mouse lines from Goesan-gun (n = 6) using Micro-CT and ImageJ: head and body (nose to 3rd coccygeal vertebra), tail (tip to 3rd coccygeal vertebra) (Fig 2), tail ratio (head and body length / tail length × 100), and cranial linear measurements (nasal, frontal, parietal, interparietal, interparietal bone, maxillary and mandible bone) (Fig 3). The craniometric variables were log-transformed to meet the assumption of normality. We compared 30 digitized landmarks in the cranium of our inbred mouse lines (n = 6, 9 weeks old) with 4 laboratory mouse strains (CBA, C57BL/6H, C3H, BALB/c) (n = 6 each, 9 weeks old). This analysis aimed to depict the morphospace of the cranium, originating from wild mice from Geosan-gun (Fig 4). Position, orientation, and scaling biases were standardized using generalized Procrustes analysis. Variations in skull shape and inter-subspecies distance between their mean shapes were examined using the geomorph package (v. 3.0.1) in R on a scatter plot of relative coordinates. All statistical analyses were done using R software (version 4.2.2).

**Fig 2.**
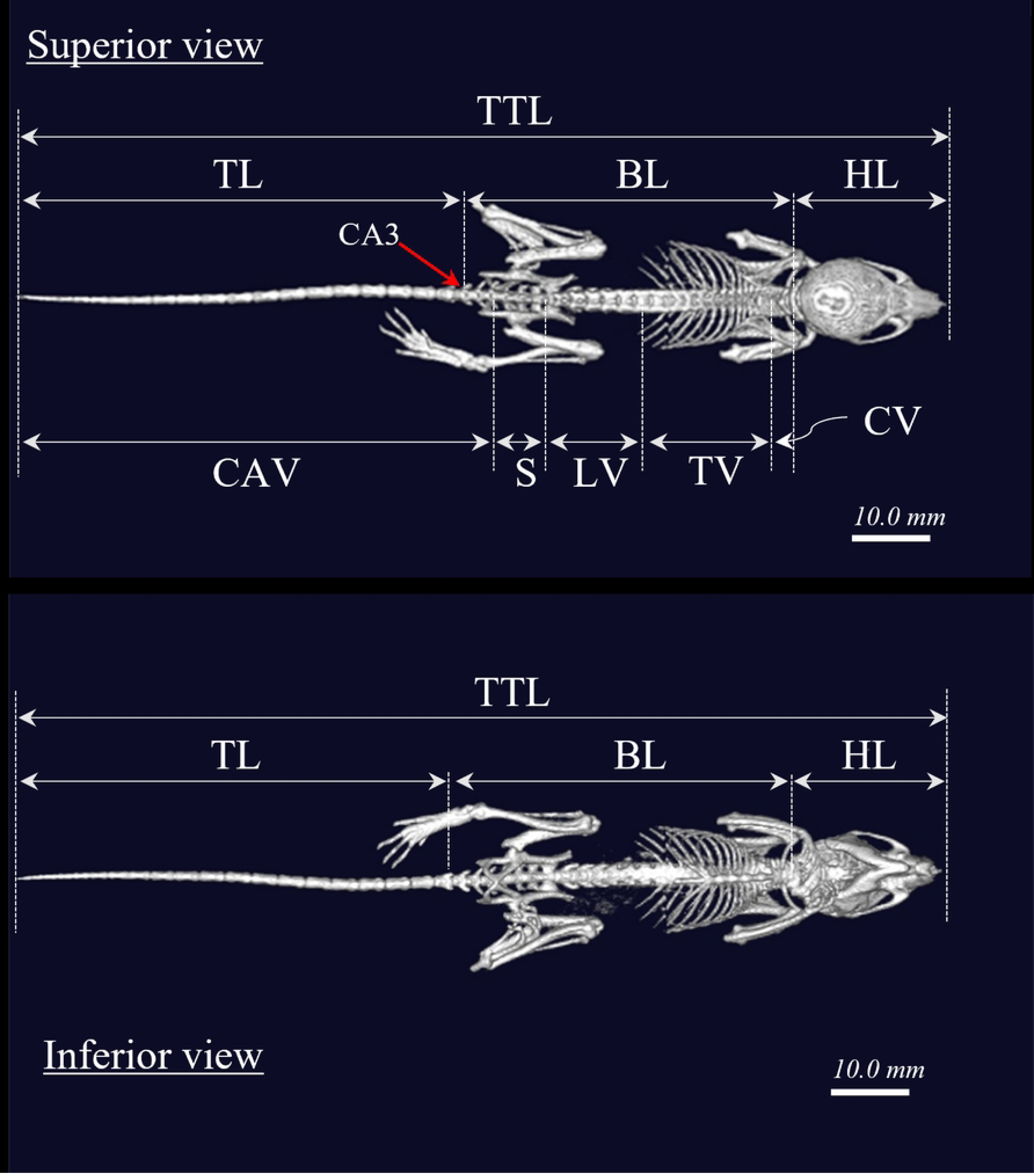
External measurements of bone length of a mouse on micro-CT images. TTL, total length; TL, tail length from tip to the no. 3 coccygeal vertebra; HL, head length; BL, body length; HL + BL, lengths of the head and body determined from the nose to no. 3 coccygeal vertebra; Tail ratio: HL + BL/ TL × 100); CA3, no. 3 coccygeal vertebra; CV, cervical vertebrae; TV, thoracic vertebrae; LV, lumbar vertebrae; S, sacrum; CAV, caudal vertebrae.

**Fig 3.**
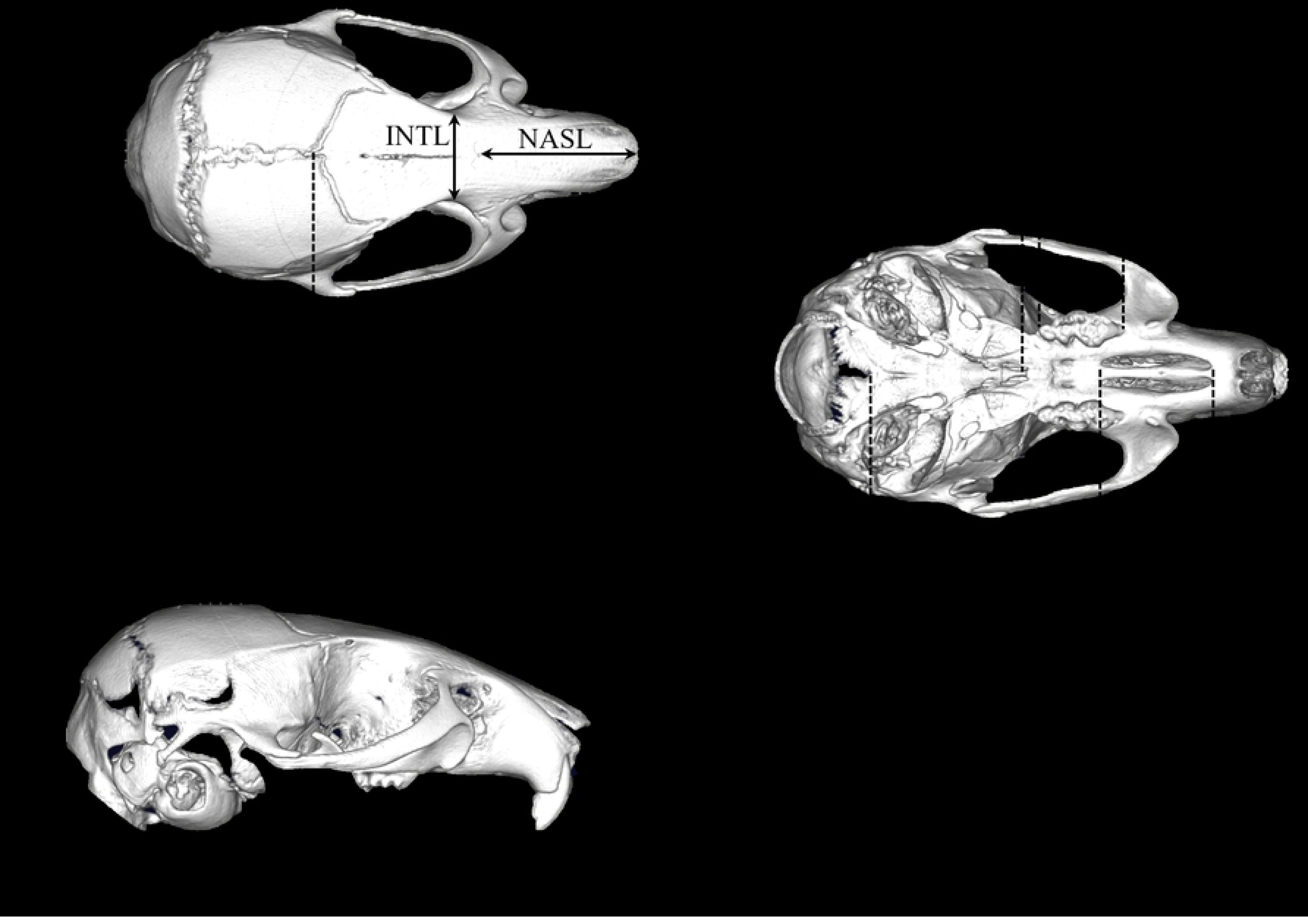
Linear measurements of the cranial and dental variables on micro-CT images. Abbreviations for the 14 cranial variables are as follows: OccL, occipitonasal length; ZYGW, maximum zygomatic arches’ width; INTL, interorbital constriction width; NASL, nasal length; PMXW, premaxillary width; FNL, frontonasal length; BraH, braincase height on tympanic bullae; FL, frontal bone length; MXTL, maxillary teeth row length; ForL, foramina incisive length; CBL, condylobasal length; BASL, basal length; BSIL, basilar length; NW: nasal bone width.

**Fig 4.**
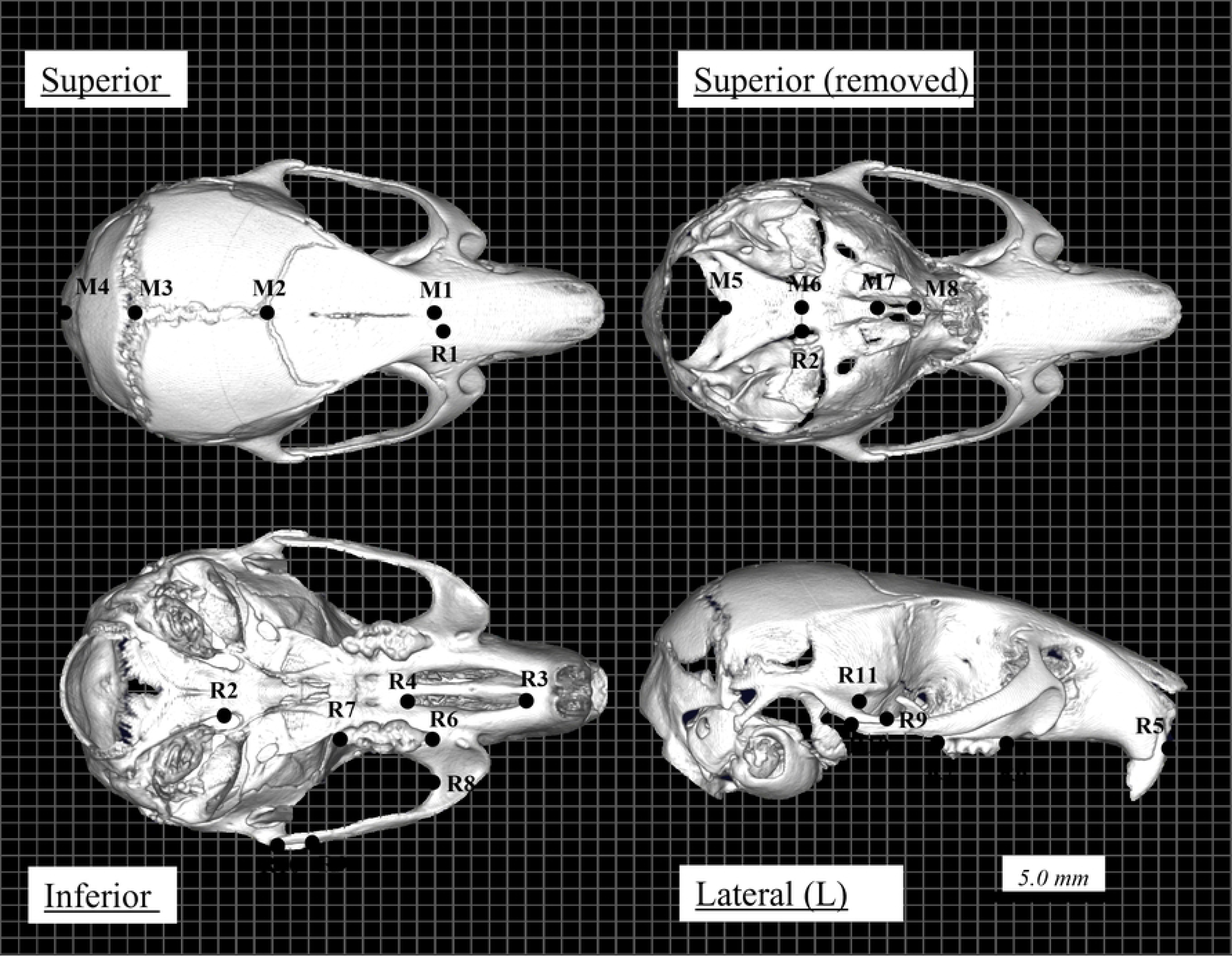
Digitized landmarks (M1 ∼ 8; R1 ∼ 11) in the cranium. M1, posterior of nasal bone; M2, intersection of coronals suture and sagittal suture; M3, intersection of anterior lambdoid suture and sagittal suture; M4, caudal of skull; M5, lowest point of foramen magnum; M6, intersection of basisphenoid occipital bone; M7, intersection of presphenoid basisphenoid; M8, anterior of presphenoid; R1, intersection of premaxilla, nasal bone and front bone; R2, intersection of basisphenoid occipital bone; R3, rostral of rostral palatine foramen; R4, caudal of rostral palatine foramen; R5, frontal tip of incisors; R6, rostral tip of 1st molar; R7, caudal tip of 3rd molar; R8, lowest point of zygomatic bone; R9, rostral point of zygomaticotemporal suture; R10, caudal of zygomaticotemporal suture; R11, posterior tip of zygomatic arch.

## Results

### Phylogeny among Korean mice and their relationships to the *M. musculus* subspecies

We combined the 23 *cytb* (1,140bp) sequences with publicly accessible 169 data set of *M. musculus* subspecies from Korea and other countries (S1 Table). The ML phylogeny from *cytb* data set splits into three discrete clades corresponding to the CAS, DOM, and MUS with BS/PP values >95% (Fig 5a), a well-differentiated cluster of MUS, which includes all the Korean mice (Figs 5a and 5b). Although previously designated as a distinct subspecies from hybrids between MUS and CAS in Japan, *M. m. molossinus* (MOL) segregates within the MUS cluster. The topology of the median joining network diagram that shows optimal connections between the haplotypes of *M. musculus* subspecies also support three discrete subspecies clusters (Fig 6a). The Korean mice from the network of *cytb* haplotypes are confined within the MUS populations with unique haplotypes (e.g., *Hap*_1 ∼ 3 and *Hap*_5 ∼ 6), sharing some haplotypes in basal positions with Chinese populations (e.g., *Hap*_4 and *Hap*_7) (Fig 6b, S1 Table). The shared haplotypes in basal positions from Japan with Korean (e.g., *Hap*_7 ∼ 8) and Chinese (e.g., *Hap*_7) populations (Fig 6b, S1 Table) further supports the close relationship between population expansion, rendering a geographical distribution stretching from northern China over the Korean Peninsula into Japan. Additionally, MOL holds 4 haplotypes (Fig 6b, S1 Table), where it shares a haplotype (e.g, *Hap*_7) with MUS among China, Korea, and Japan, consistent with their clustering within the MUS sub-lineage (Fig 5a).

**Fig 5.**
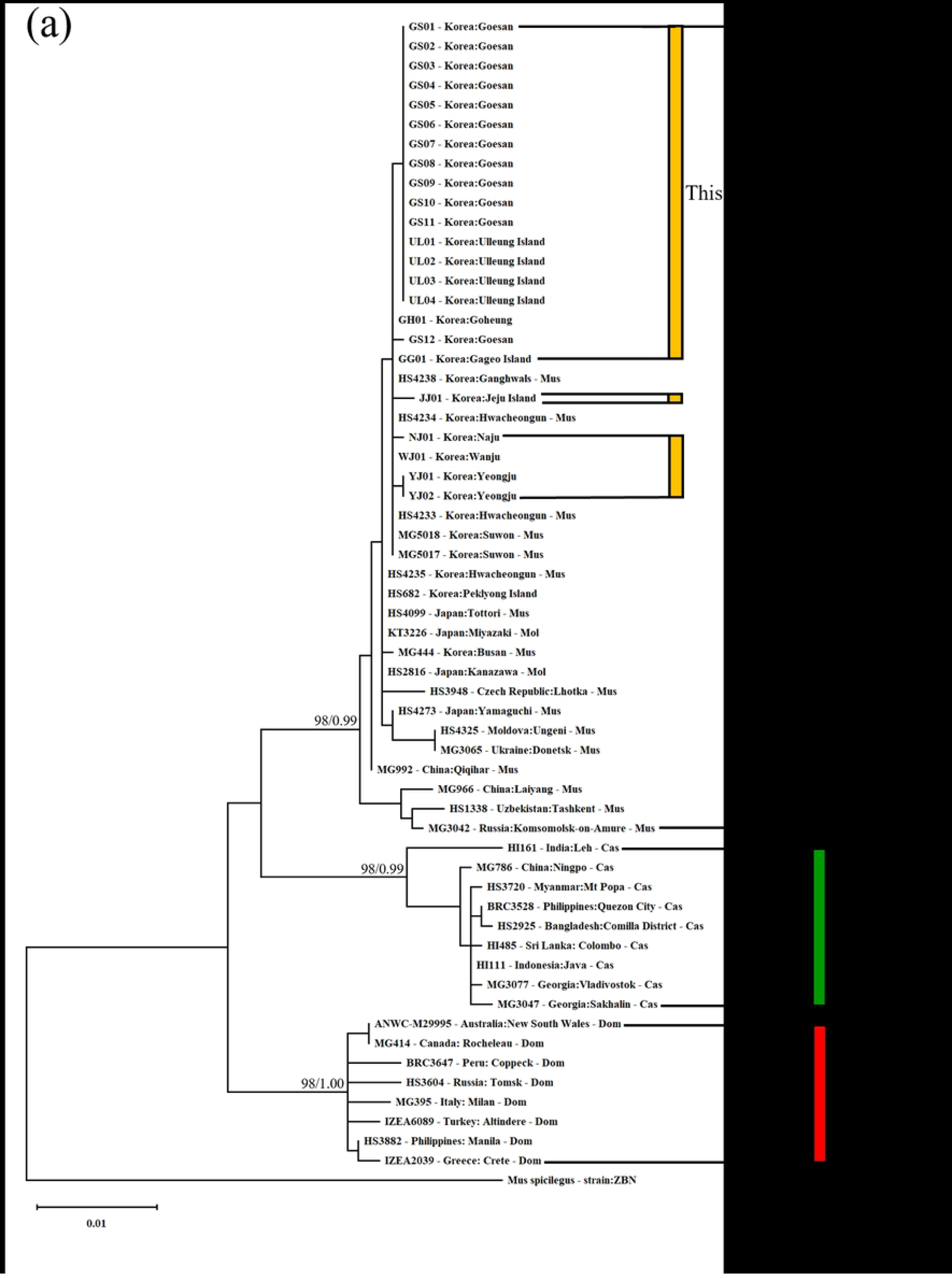
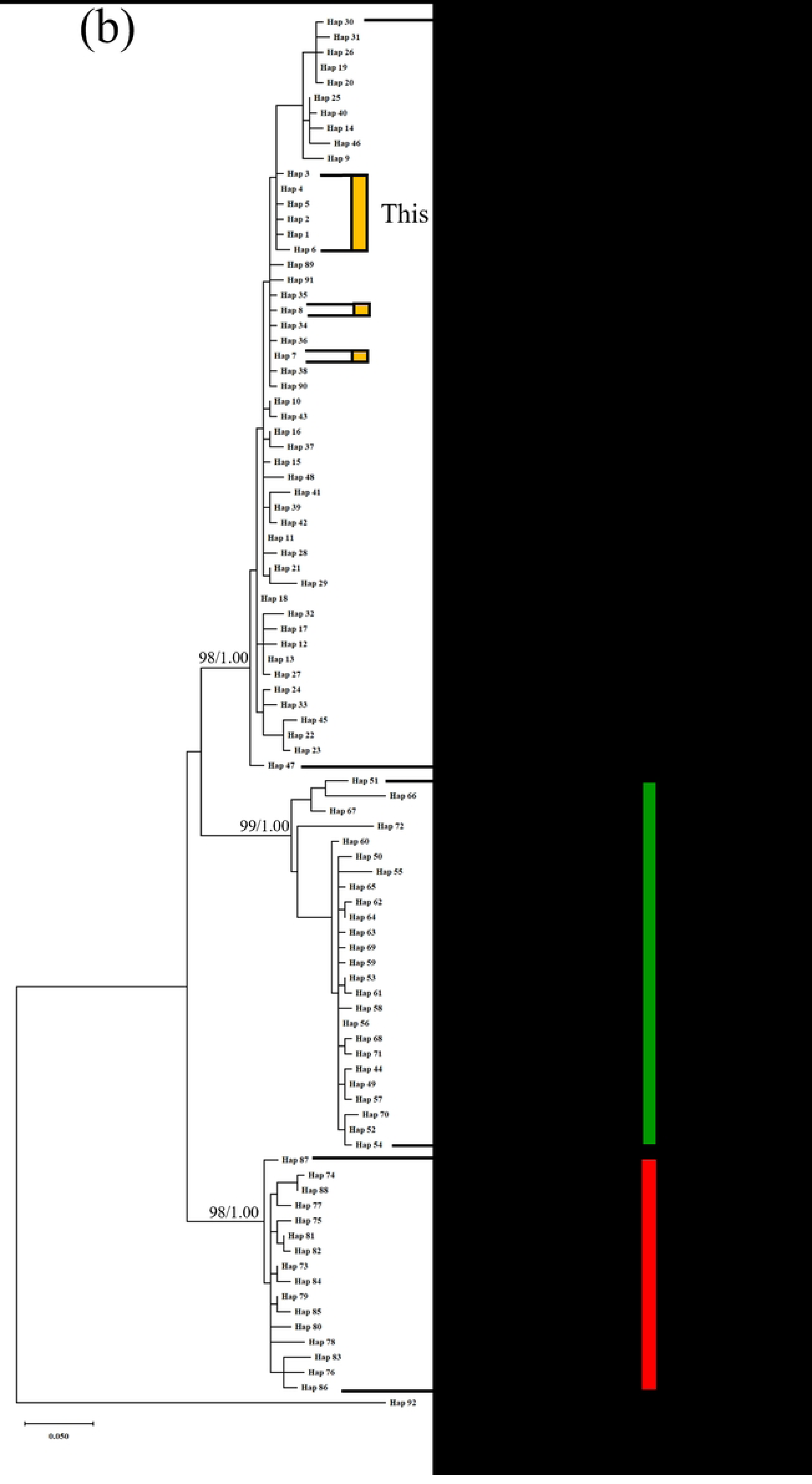
Maximum likelihood (ML) and Bayesian Inference (BI) phylogenies based on mitochondrial *cytb* sequences of *Mus musculus* (a). Reconstructed ML/BI tree from haplotype of mitochondrial *cytb* sequences of *Mus musculus* (b). The phylogroups represent the three subspecies groups: *M. m. musculus* (MUS), *M. m. castaneus* (CAS), and *M. m. domesticus* (DOM). The evolutionary history was inferred by using the ML/BI method and Hasegawa-Kishino-Yano model with the *cytb* data sets (S1 Table). ML bootstrap/BI posterior probability (BS/PP) support values (1000 replicates) are indicated above the nodes. *Mus spicilegus* (strain: ZBN) is used as an outgroup.

**Fig 6.**
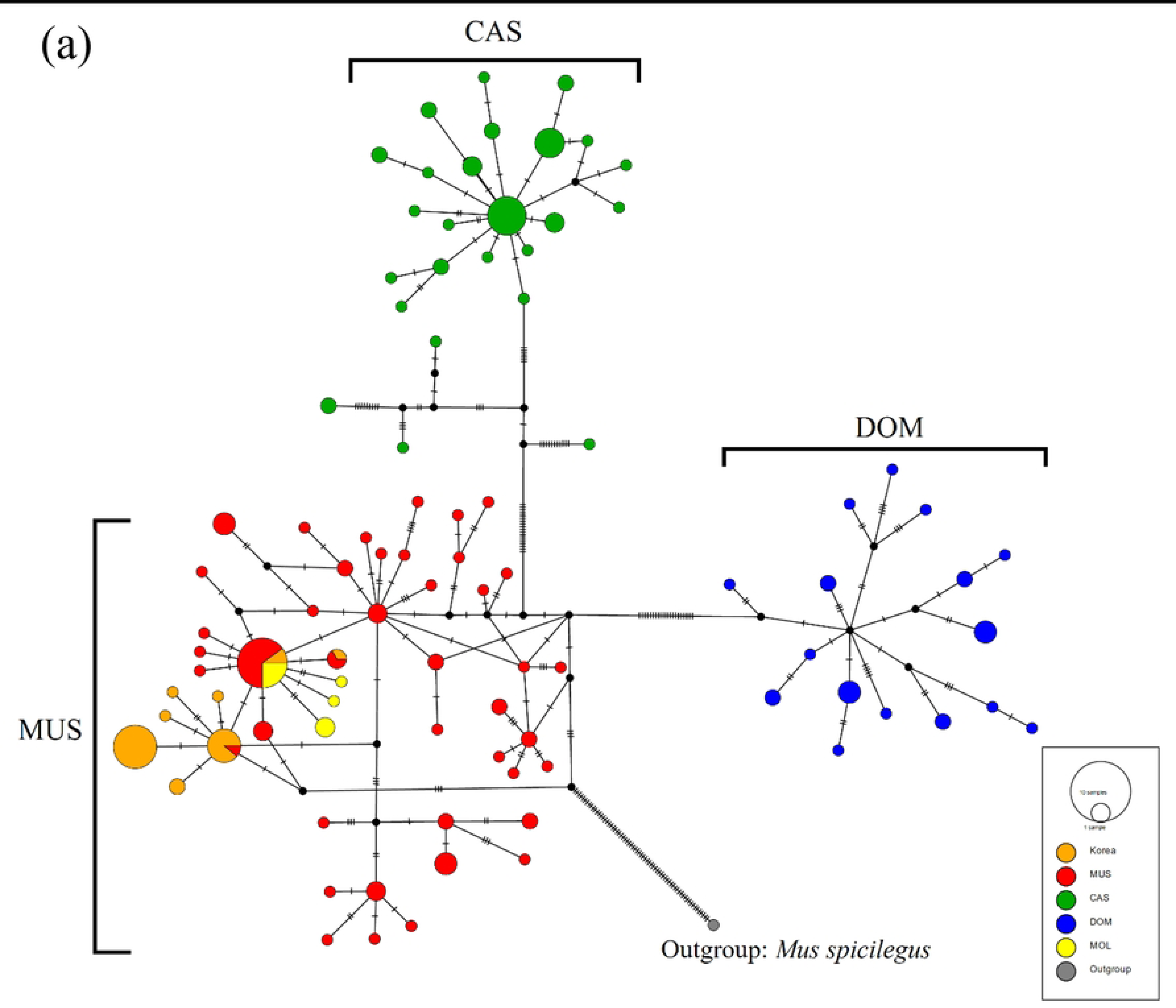
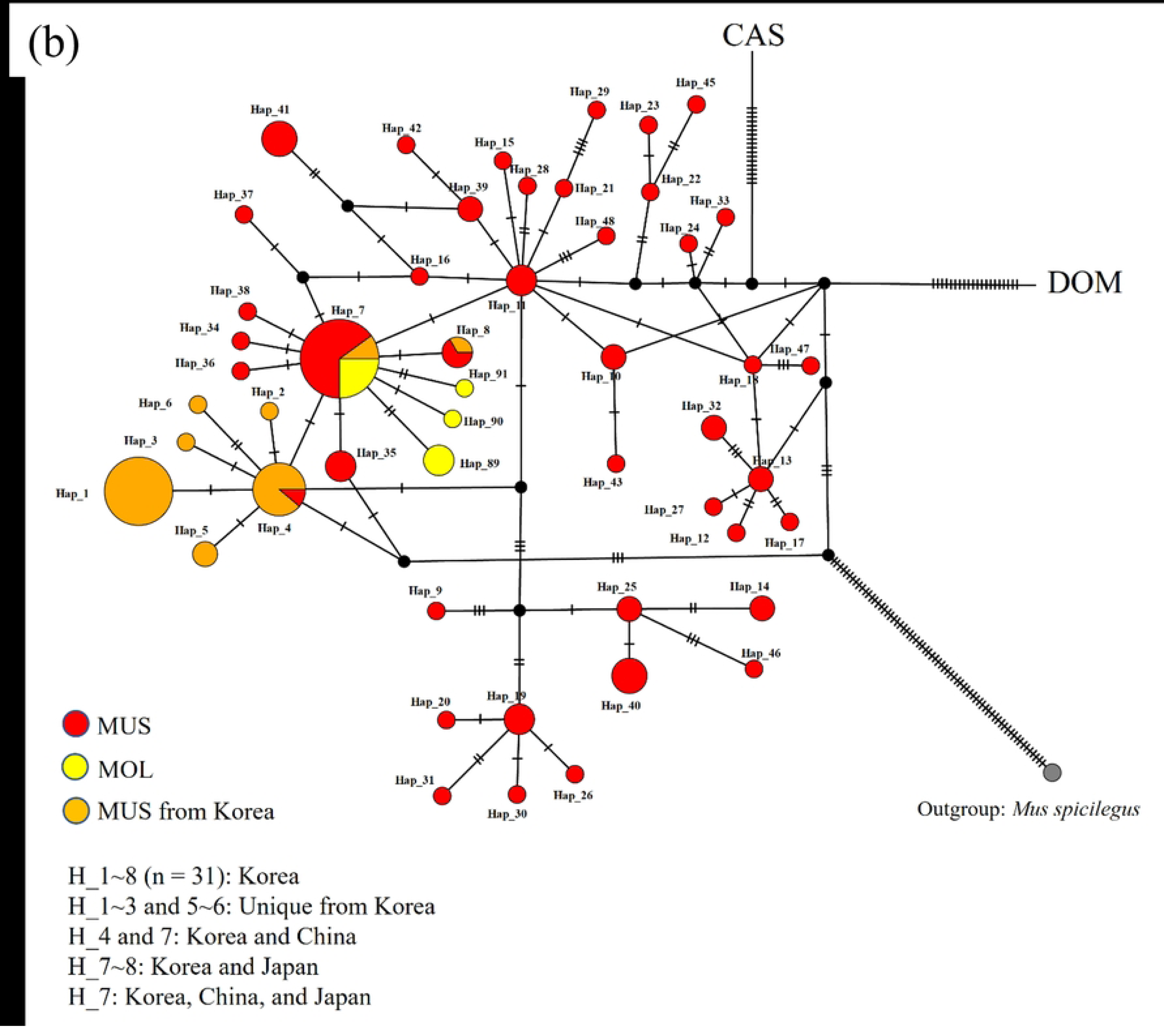
Haplotype networks tree based on the mitochondrial *cytb* sequences of *Mus musculus.* **(a).** The phylogroups represent the three subspecies groups: *M. m. musculus* (MUS), *M. m. castaneus* (CAS), and *M. m. domesticus* (DOM). We retrieved publicly available *cytb* sequences belonging to *M. musculus subspecies* (S1 Table), along with representative sequences of closely related species as an outgroup (e.g., *Mus spicilegus* – strain: ZBN). **Haplotype networks tree for MUS and MOL (*M. m. molossinus*) populations based on the mitochondrial *cytb* sequences of *Mus musculus* (b).** Pairwise genetic distances (*p*-distance) between Korean mice and recognized subspecies populations ranged from 0.46% and 2.94%. Korean mice are closest to the MUS lineage, including MOL from Japan, whereas they are most distantly related to CAS, showing the highest genetic divergence (Table 1). *Fst* values also support pairwise genetic differentiation among and within subspecific populations, indicating significant evolutionary differentiation between the three primary subspecies (Table 2). Genetic variations do not differ significantly between MUS populations and Korean mice (*Fst* = 0.188, *p* > 0.05). However, Korean mice within the MUS group remain distinct from CAS (*Fst* = 0.902, *p* < 0.0001) and DOM populations (*Fst* = 0.889, *p* < 0.0001), although not significanlty different from MOL (*Fst* = 0.490, *p* > 0.05) (Table 2).

**Table 1.**
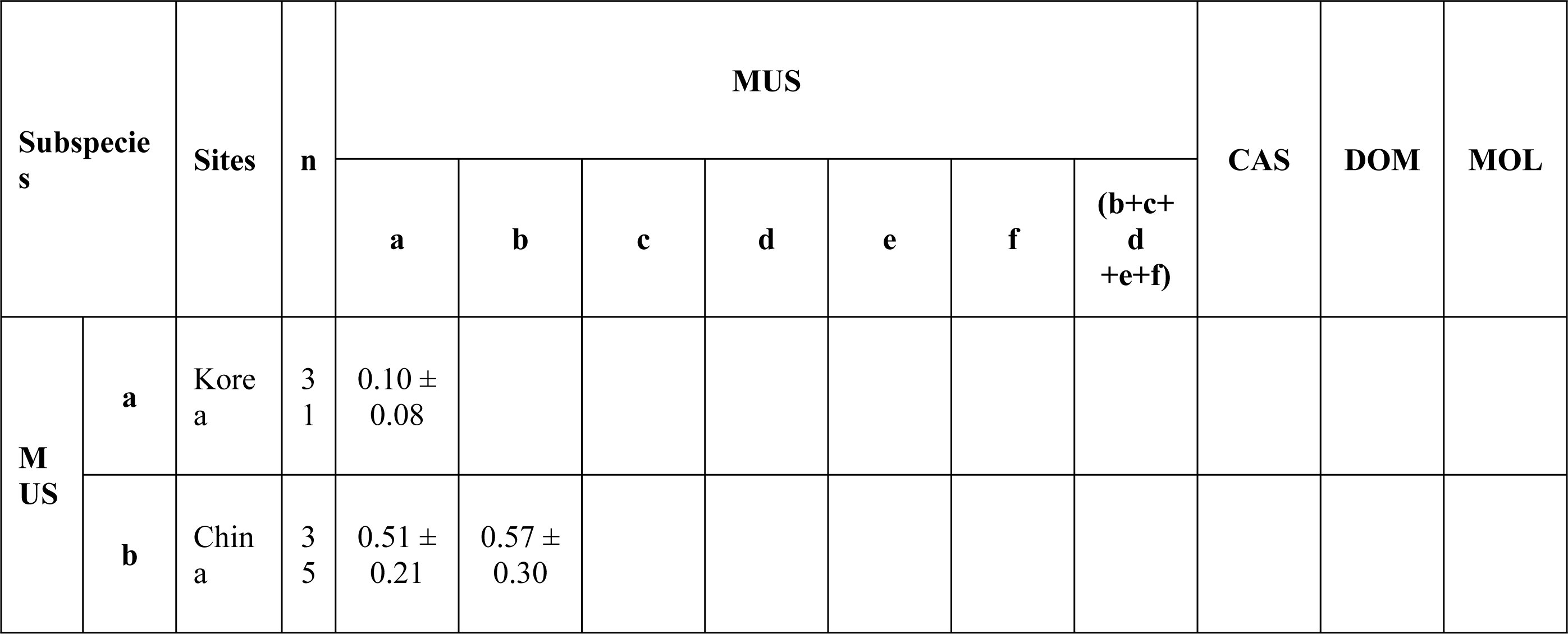
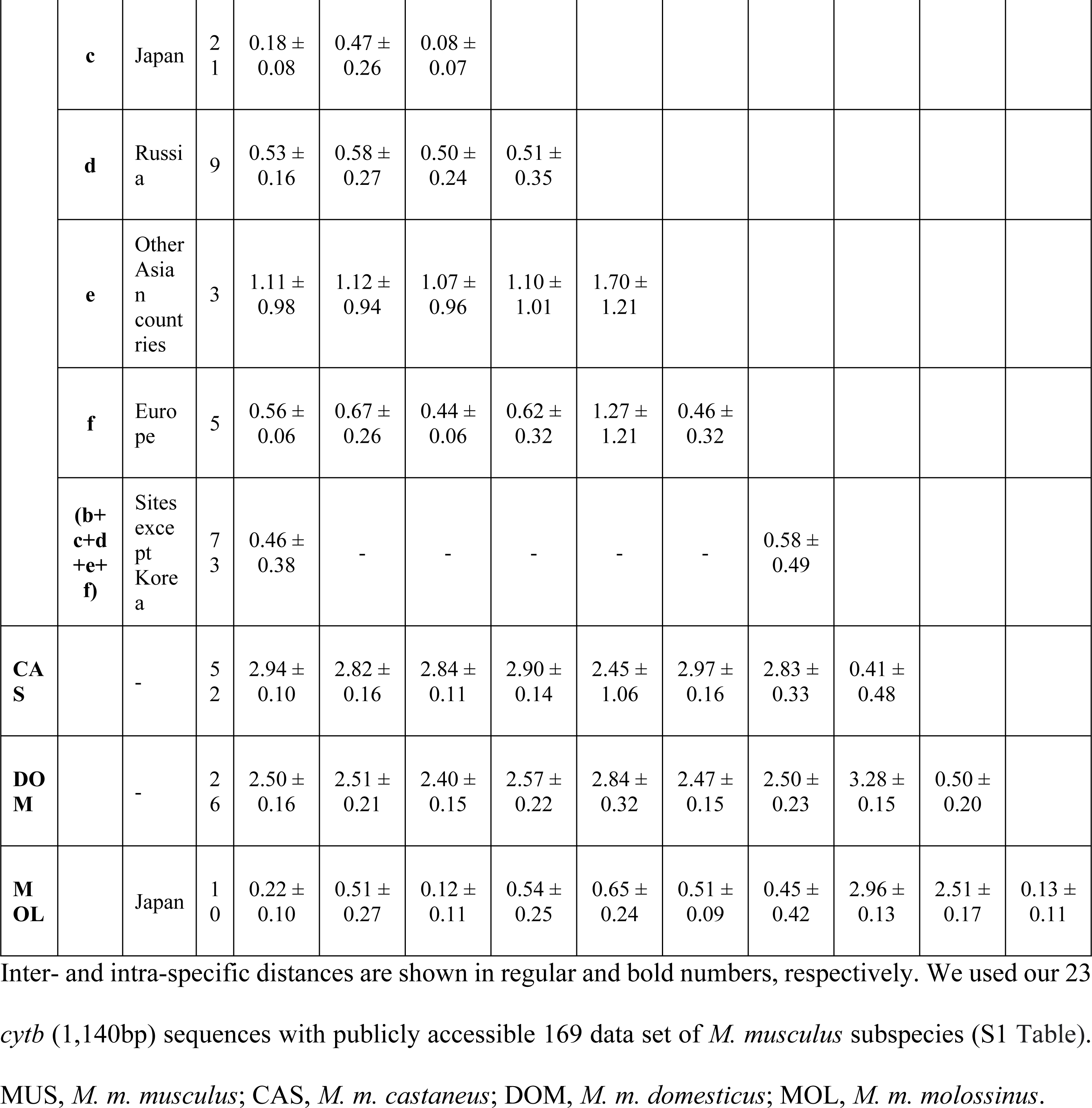
Genetic distance (*p*-distance ± SD; %) within and between *Mus musculus* subspecies based on mitochondrial *cytb* sequences.

**Table 2.**
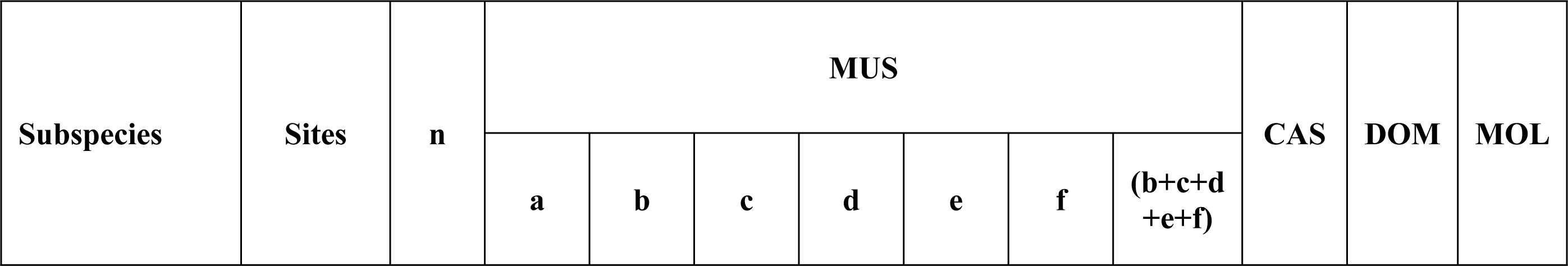
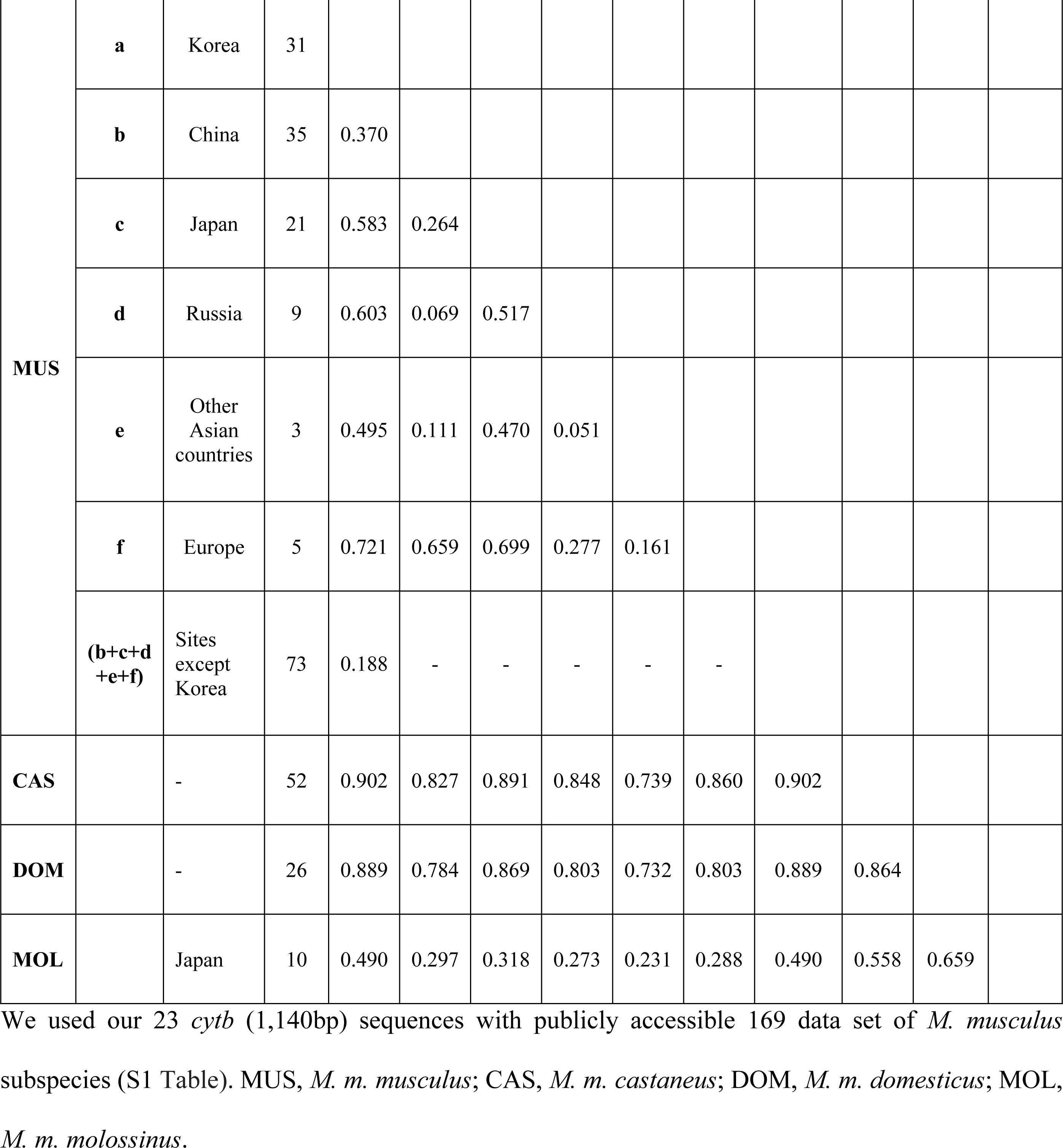
The pairwise *Fst* values for genetic differentiation among and within subspecific populations based on mitochondrial *cytb* sequences of *Mus musculus*.

Genetic diversity and demographic statistics are summarized in Table 3. The G+C content of the *cytb* sequences is almost identical across *M. musculus* populations, ranging from 38.9% to 39.1%. Haplotype diversity (*Hd*) in each major subspecies ranged from 0.925 to 0.953, followed by MOL (0.711) and the lowest in the Korean population (0.071). The nucleotide diversity (*π*) ranged from 0.44% in MUS to 0.51% in DOM, with the lowest in the Korean mice (0.10%). Tajima’s *D* values, a statistical test for neutrality of mutations, were not significant for most sample locations except Southeast Asia. However, negative values in pooled samples from both MUS (−1.9628, *p* < 0.05) and CAS (−2.0617, *p* < 0.05) suggest a recent population expansion.

**Table 3.** Genetic diversity and demographic statistics of *Mus musculus* subspecies based on mitochondrial *cytb* sequences.

| Groups | | n | $\pi$ | $H$ | $Hd$ | Tajima's $D$ | Fu's $F_s$ | G+C content |
| --- | --- | --- | --- | --- | --- | --- | --- | --- |
| Korea | | 31 | 0.0010 ± 0.0002 | 8 | 0.071 ± 0.068 | -1.2924 | -3.322 | 0.391 |
| MUS | | 73 | 0.0044 ± 0.0004 | 41 | 0.953 ± 0.016 | **-1.9628\*** | -31.286 | 0.390 |
| | China | 35 | 0.0053 ± 0.0004 | 27 | 0.983 ± 0.011 | -1.4668 | -17.605 | 0.390 |
| | Japan | 21 | 0.0008 ± 0.0002 | 7 | 0.667 ± 0.107 | -1.7186 | -3.780 | 0.391 |
| | Russia | 9 | 0.0051 ± 0.0007 | 5 | 0.806 ± 0.120 | 0.5839 | 1.557 | 0.390 |
| | Other Asian countries | 3 | 0.0170 ± 0.0058 | 5 | 1.000 ± 0.126 | -0.7674 | 0.485 | 0.389 |
| | Europe | 5 | 0.0046 ± 0.0014 | 3 | 0.700 ± 0.218 | -0.1091 | 2.385 | 0.391 |
| CAS | | 52 | 0.0049 ± 0.0010 | 25 | 0.925 ± 0.024 | **-2.0617\*** | -10.853 | 0.387 |
| | China | 14 | 0.0022 ± 0.0008 | 5 | 0.857 ± 0.108 | -1.3200 | -0.640 | 0.388 |
|  | Japan | 2 | - | 1 | - | - | - | 0.388 |
| | India | 12 | 0.0094 ± 0.0021 | 8 | 0.945 ± 0.054 | -0.7320 | 0.410 | 0.389 |
| | Southeast Asia | 20 | 0.0023 ± 0.0008 | 11 | 0.868 ± 0.064 | **-2.0438\*** | -4.605 | 0.388 |
| | Europe | 4 | 0.0013 ± 0.0007 | 2 | 0.500 ± 0.265 | -0.7545 | 1.716 | 0.388 |
| DOM | | 26 | 0.0051 ± 0.0004 | 16 | 0.951 ± 0.024 | -1.6938 | -4.478 | 0.390 |
| | Russia | 4 | 0.0061 ± 0.0017 | 3 | 0.833 ± 0.222 | -0.1297 | 2.038 | 0.390 |
| | Asia and Northern Africa | 3 | 0.0041 ± 0.0012 | 3 | - | - | 0.308 | 0.390 |
| | Europe | 15 | 0.0052 ± 0.0005 | 12 | 0.971 ± 0.033 | -1.5965 | -3.999 | 0.390 |
|  | **Oceania** | 2 | - | 1 | - | - | - | 0.390 |
|  | **America** | 2 | 0.0061 ± 0.0031 | 2 | - | - | - | 0.392 |
| **MOL** | **Japan** | 10 | 0.0014 ± 0.0003 | 4 | 0.711 ± 0.117 | -0.5311 | 0.017 | 0.391 |

We used our 23 *cytb* (1,140bp) sequences with publicly accessible 169 data set of *M. musculus* subspecies (S1 Table). n, number of samples; $\pi$ , nucleotide diversity; *H*, number of haplotype; *Hd*, haplotype diversity, MUS, *M. m. musculus*; CAS, *M. m. castaneus*; DOM, *M. m. domesticus*; MOL, *M. m. molossinus*. Significant demographic parameters are indicated in bold with asterisk (\* $p < 0.05$ ).

### Morphometrics

Considering the haplotypes of the present Korean mice as MUS, Fig 7 and Table 4 present the tail ratios from our samples and previously described subspecies populations. The tail ratios of our Korean mice (ranging from 104.8 ∼ 140.0%) were similar to those previously reported for Korean mice assigned to the subspecies *M. m. molossinus* (107.7%) [2], *M. m. utsuryonis* (105.1%) [4], and *M. m. yamashinai* (121.7%) [5]. The CAS subspecies is characterized by a tail length exceeding the head and body length [2]. This contradicts a recent molecular study showing tail ratios above 100% for CAS [17]. In contrast, the typical tail length of MUS and MOL is shorter than the head and body length [2], which aligns with all our observations, including the Korean mice (Fig 7).

**Fig 7.**
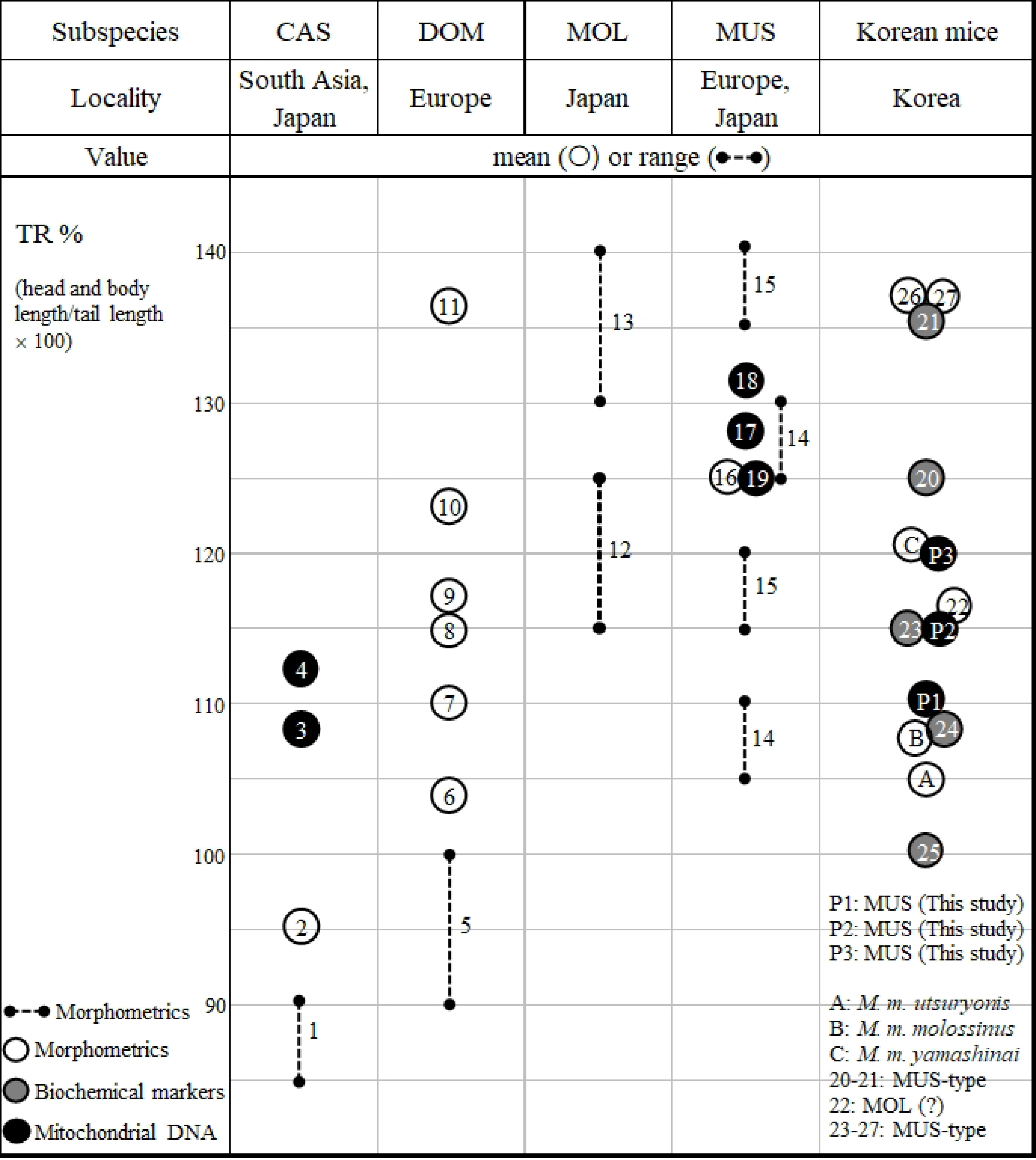
Comparison of tail ratios (TR%) from the present Korean mice and formerly characterized subspecies populations. MUS, *M. m. musculus*; CAS, *M. m. castaneus*; DOM, *M. m. domesticus*; MOL, *M. m. molossinus*. Numbers in the figure indicate data from the references as shown in Table 4: P1-P3 [this study], (A) [4], (B) [3], (C) [5], (1, 5, and 7-15) [2], (2, 6, and 16) [8], (3, 4, and 19) [17], (17 and 18) [18], (20 and 21) [14], (22) [9], (23-25) [10], (26 and 27) [11].

**Table 4.** Tail ratios (head and body length/tail length × 100) of wild house mice from the present study and formerly characterized subspecies populations.

|  | Method | Locality | Type locality | Tail ratio (%) |  | Remarks | Reference |
| --- | --- | --- | --- | --- | --- | --- | --- |
|  |  |  |  | Mean | Range |  |  |
| MUS | a | Sweden, Esthonia | Upsala, Sweden |  | 115 ~ 120 | Variation: males, 92 ~ 143; females, 97 ~ 155 | [2] |
| (Linnaeus, 1758) |  |  |  |  | 135 ~ 140 |  |  |
|  | a | East Prussia, Hungary |  |  | 105 ~ 110 |  |  |
|  |  | **Korea** |  |  |  |  |  |
| *M. m. yamashinai* | a | Mokpo and its neighboring islands, and inlands | Mokpo, Korea | 121.7 (n = 10) | 103.5 ~ 136.2 | Originally *M. Bactrianus yamashinai* | [5] |
| *M. m. molossinus* | a | Jeju-do (islands) | - | 107.7 |  |  | [2,3] |
| *M. m. utsuryonis* | a | Ulleung-do (islands) | Ulleung, Korea | 105.1 (n = 5) | 98.4 ~ 112.3 | Originally *M. molossinus utsuryonis* | [4] |
| MUS | b | Geosan |  | 111.1 (n = 6) | 109.0 ~ 113.7 | Rice field and granaries | [This study] |
| MUS | b | Yeongju, Wanjū |  | 120.8 (n = 3) | 105.1 ~ 140.0 | Mountain areas |  |
| MUS | b | Ulleung-do, Gageo-do |  | 115.4 (n = 4) | 104.8 ~ 128.0 | Islands |  |
| MUS-like | c | Chuncheon |  | 125.6 (male) |  | Based on AKV-type murine leukemia virus gene | [14] |
|  | c |  |  | 136 (female) |  |  |  |
| - | a | Chuncheon |  | 116.0 |  | Consistent with *M. m. yamashinai* | [9] |
| MUS-like | c | Chuncheon |  | 115.1 |  |  | [10] |
| MUS-like | c | Sokcho |  | 108.4 |  |  |  |
| MUS-like | c | Heongsung |  | 100.1 |  |  |  |
| MUS-like | a | Anmyun-do (islands) |  | 137.1 |  |  | [11] |
| MUS-like | a | Gumi |  | 137.6 |  |  |  |
| MUS | b | Honshu, Japan |  | 125.0 |  | *cytb* | [17] |
| MUS | b | Miura, Japan |  | 132.2 |  | *cytb* | [18] |
| MUS | b | Fujisawa, Japan |  | 127.5 |  | *cytb* |  |
| MUS | a | Northern Europe |  | 125.3 |  |  | [8] |
| CAS | a | South Asia | Philippine Islands |  | 85 ~ 90 |  | [2] |
| (Waterhouse, 1843) | b | Hokkaido, Japan |  | 108.4 |  | *cytb* | [17] |
|  | b | Honshu, Japan |  | 112.4 |  | *cytb* |  |
|  | a | Southeastern Asia |  | 95.6 |  |  | [8] |
| DOM | a | Switzerland | Dublin, Ireland | 115 |  |  | [2] |
| (Rutty, 1772) | a | France |  | 110 |  |  |  |
|  | a | Ireland |  | 123 |  |  |  |
|  | a | Shetland Islands |  | 117 |  |  |  |
|  | a | St. Kilda |  | 136 |  |  |  |
|  | a | somewhere |  |  | 90 ~ 100 |  |  |
|  | a | Europe |  | 104.0 |  |  | [8] |
| MOL | a | Japan | Nagasaki/Kyushu, Japan |  | 115 ~ 125 (male) | Variation: males, 119 ~ 138; females, 120 ~ 162 | [2] |
| (Temminck, 1845) | a |  |  |  | 130 ~ 140 (female) |  |  |

a (morphometrics), b (mitochondrial DNA), c (biochemical markers). MUS, *M. m. musculus*; CAS, *M. m. castaneus*; DOM, *M. m. domesticus*; MOL, *M. m. molossinus*.

We measured linear distances between specific landmark points on the cranium (boundaries between different skull bones) from 3D micro-CT images to characterize the craniometric structure of Korean mice (Figs 3 and 4). Except for maxillary tooth-row length (MXTL), all measured variables (PMXW, INTL, ZYGW, NASL, FNL, BSIL, BASL, BULL, CBL, ForL, DIAL, ItoM, FL, and BraH) were significantly shorter in Korean mice compared to the four mouse strains (CBA, C57BL/6H, C3H, and BALB/c) (*p* < 0.05), with no significant sex differences (Data not shown). To account for size differences between populations with 9-week-old mice (only adult specimens with skull CBL >17.0 mm [19]), we used standardized craniometric indices. These indices are ratios relative to CBL and were log-transformed for analysis (Fig 8). The PMXW (premaxillary tooth-patch width) and MXTL values indicate that Korean mice have a narrower nasal bone and larger molars compared to the other strains derived from DOM (*p* < 0.05) (Fig 8). Other variables were not statistically different, except for basilar length (BSIL), which varied among populations with a significant difference between Korean mice and CH3 strains (*p* < 0.05) (Fig 8). Despite not being statistically significant, log-transformed craniofacial measurements of FNL, CBL, NASL, FL, and BASL in our Korean MUS mice appear smaller than those from several subspecies, including DOM, CAS (previously assigned *M. m. bactrianus*, now recognized as genetically indistinguishable from CAS [20], MUS, and secondary populations (e.g., *M. m. isatissus*) from the Middle East [21] (S1 Fig). External measurements of NW, NASL, and FNL are likely shorter in our Korean MUS and MOL (MSM/Ms) compared to other inbred mouse lines derived from DOM (BALB/c, C3H, C57BL/6H, CBA, ICR). However, the relative ratios between NW and NASL were comparable among these groups [22] (S2 Fig). These highly heritable craniometric features indicate that differences between subspecies are more distinct than variations within subspecies, although genetically close strains do not always possess morphologically identical crania.

**Fig 8.**
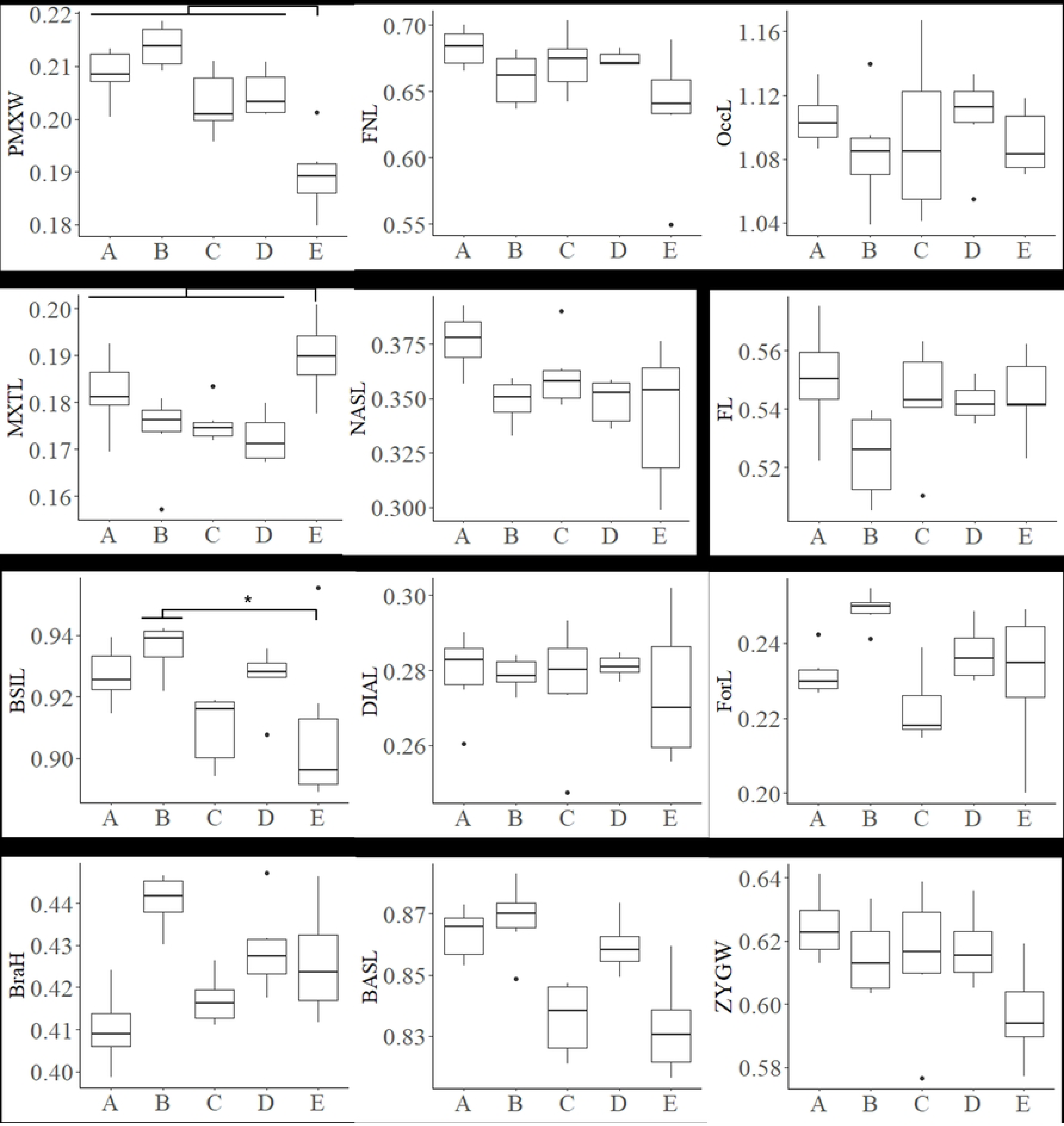
Box plot of craniometric variables with statistically significant level among 4 commercially available laboratory mouse strains (CBA, C57BL/6H, C3H, and BALB/c derived from DOM) and our inbred mice from Goesan-gun (KG01BRA/1CNUf1∼f3 and KG01BRA/1CNUm1∼m3). No sex difference in each variable (3 males and 3 females each with 9 weeks old) was found in all the strains including our inbred mice, thus all mice with only adult specimens (n = 6 each; >17.0 mm of the condylobasal length (CBL) of the skull [19]) were further combined. Box plot shows mean, range, and minimum and maximum values in the figure. A, CBA; B, C57BL/6H; C, C3H; D, BALB/c; E, our inbred mice from Goesan-gun. To account for size differences between populations with only adult specimens (Y-axis), we used standardized craniometric indices. These indices are ratios relative to CBL and were log-transformed for analysis.

Morphospaces are visual representations of how craniometric variables define cranial shapes in space (Fig 9). The 30 digitized landmarks on the cranium were compared to reveal craniofacial features between our specimens and four laboratory strains (Fig 9). To account for the distinctive PMXW measurement, landmark coordinates in the nasal region and surrounding frontal bones were further adjusted to achieve a good fit for all specimens. This reflects the Korean mouse having a relatively short and slender snout with a pointed tip. The maxillary contours also indicate that the tips of the alveolar processes (tooth sockets) are positioned slightly posteriorly compared to the more elaborate configuration in DOM-derived laboratory strains. These are subtle differences between the Korean mice and the other groups.

**Fig 9.**
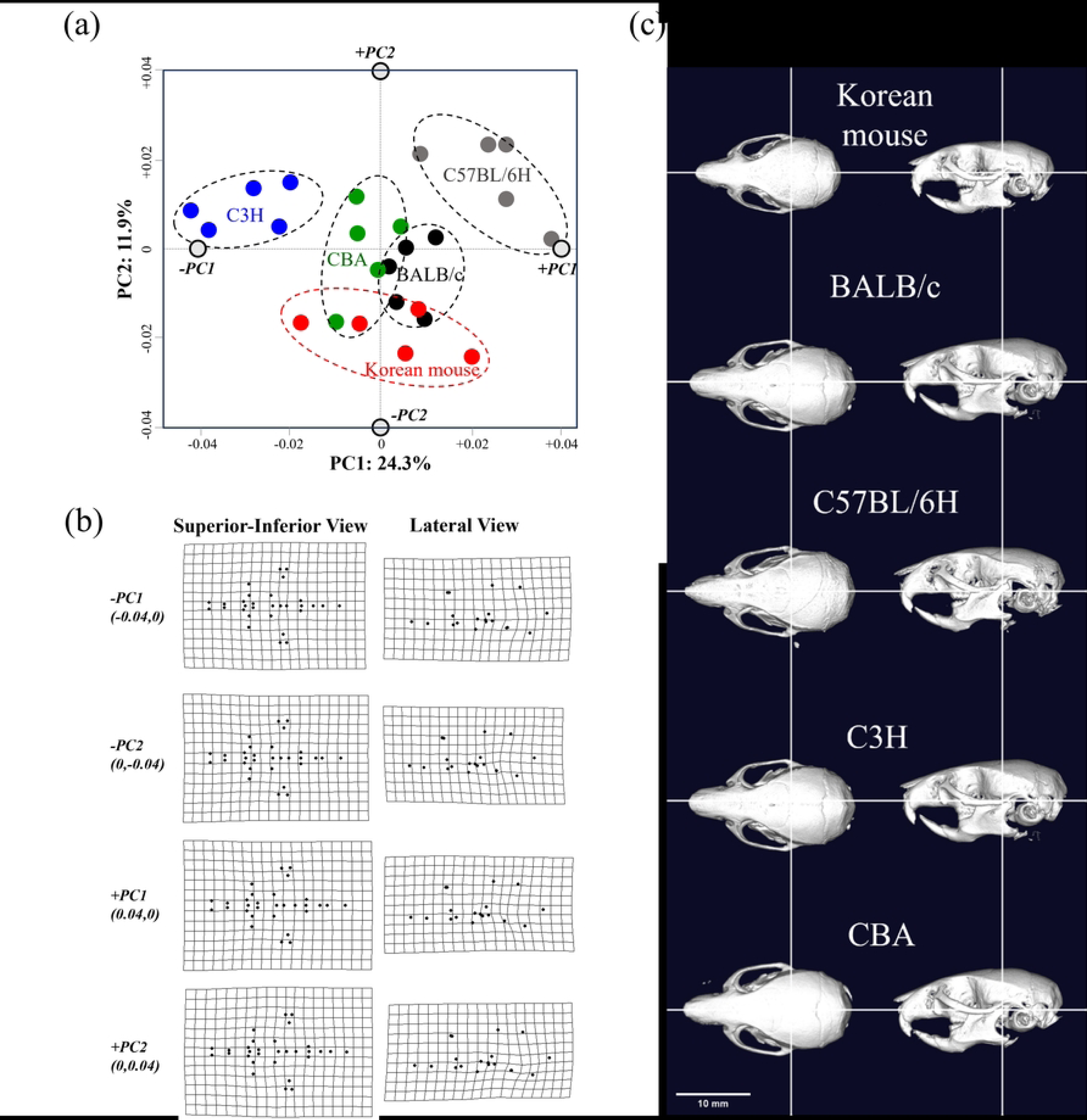
Analyses of cranial morphospaces among 4 laboratory mouse strains (CBA, C57BL/6H, C3H, and BALB/c derived from DOM) and our inbred mice from Goesan-gun (KG01BRA/1CNU). No sex difference in each variable (3 males and 2 females each with 9 weeks old) was found in all the strains including our inbred mice (n = 5 each). a) PCA analysis based on relative distance from the 30 digitized landmarks in the cranium; b) scatter plot on relative coordinates after configuration of the 30 digitized landmarks in the cranium; c) micro-CT images from the view of superior and calotte removed superior crania among 4 laboratory mouse strains and our inbred mouse from Goesan-gun.

## Discussion

Our key findings are: 1) Korean house mice represent populations of MUS with significantly lower genetic variation compared to other primary subspecies. 2) The genetic structures, including *cytb* haplotype networks, suggest close phylogeographic relationships between Korea and neighboring countries, influencing MUS population dynamics. 3) We found no evidence supporting a unique subspecies designation for mice previously assigned to *M. m. molossinus*, *M. m. utsuryonis*, and *M. m. yamashinai*. These mice likely belong to existing MUS populations. Our findings support previous studies suggesting that the CAS subspecies found in Japan migrated directly from Southeast Asia to Japan without passing through Korea. However, further studies involving additional sampling and analysis from the southern regions of the Korean Peninsula and islands, including Jeju Island, are deemed necessary to robustly validate this hypothesis.

The taxonomic status of Korean house mice has been a source of debate, with several subspecies assigned based on subtle morphological characteristics (e.g., MOL, *M. m. utsuryonis*, *M. m. yamashinai*, and MUS–like) or limited molecular markers (e.g., MOL–like, MUS–like, and MUS) (Fig 7 and Table 4) [23]. Our analysis of publicly available genetic datasets, including samples from Korean islands, mountains, and agricultural fields, assigned all Korean house mice, including our own samples, to MUS populations. These populations exhibited remarkably low nucleotide diversity (*π* = 0.0010) and haplotype diversity (*hd* = 0.071) compared to mice from China (*π* = 0.0053; *hd* = 0.983), Russia (*π* = 0.0051; *hd* = 0.806) and Europe (*π* = 0.0046; *hd* = 0.700) (Table 3). The relatively low genetic variations among mtDNA sequences are likely due to the presence of a single lineage in the Korean Peninsula, perhaps established by limited gene flow within geographically restricted matrilines. Whether these phylogeographic patterns are attributed to genetic drift following migration of ancient populations or to regional differentiation within an isolated resident population warrants further consideration.

Among the 42 MUS haplotypes identified in China (27), Korea (8), and Japan (7), several were shared in basal positions (e.g., *Hap*_4 and 7 between China and Korea; *Hap*_7 and 8 between Korea and Japan; *Hap*_7 among all three countries) (Fig 6b, Tables 4 and S1). This supports the hypothesis that Korean MUS originated from a single lineage in northern China, which then expanded into the Japanese Archipelago [6,7,24]. Tajima’s *D* and Fu’s *Fs* neutrality tests showed no significant deviations from neutrality for MUS haplotypes in each location, including Korea, except for Southeast Asia (Table 3), suggesting that these local populations are currently undergoing neutral evolution. However, pooling data across populations resulted in a more negative Tajima’s *D* value (−1.9628, *p* < 0.05). This could indicate a higher proportion of genetic variation within matrilines in MUS. Pooling data can be a signal of rapid population expansion, range expansion, or purifying selection events leading to population subdivision, although population structure in interaction with the sampling scheme and sample size can distort the interpretation of these neutrality tests and their reflection of regional demographic history.

While tail ratios are a valuable tool for assessing general morphological features and curtailing the influence of age-related variations in mice [8,18,19], our data show variation within and among subspecies populations (Fig 7). Therefore, tail ratios alone are insufficient for definitive subspecific classification. Previously, researchers assigned three subspecies to different Korean locations based on morphology (e.g., *M. m. molossinus* on Jeju island, *M. m. utsuryonis* on Ulleung island, and *M. m. yamashinai* on the Korean Peninsula) (Fig 1, [2–5]). This classification system has served as the official taxonomy for nearly a century [25]. Tail ratios of our wild mice (range: 104.8 ∼ 128.0%; mean: 115.4%) were comparable to previously reported values (range: 98.4 ∼ 112.3%; mean: 105.1%) from Ulleung island, suggesting they belong to the same subspecific group with some natural variation. Similarly, the tail ratios of all our specimens (range: 104.8 ∼ 140.0%) were also overlapped with those reported for *M. m. molossinus* (107.7%), *M. m. utsuryonis* (105.1%), and *M. m. yamashinai* (121.7%). Based on our phylogeographic re-evaluation, Korean mice appear to represent a single lineage within MUS. This suggests that the previously assigned subspecies (e.g., *M. m. molossinus* on Jeju island, *M. m. utsuryonis* on Ulleung island, and *M. m. yamashinai* on the Korean Peninsula) likely fall within the natural variation of MUS populations, even though genetic data for these specific subspecies is unavailable.

Genetic and/or epigenetic factors exert cranial morphology during development, making it a highly heritable trait in mice [22,26,27]. In this study, we used relative ratios of inter-landmark distances to capture the multidimensional nature of cranial morphology (Fig 9). Our 3D micro-CT-based landmark data reveal a distinct craniofacial shape in Korean mice compared to DOM-derived inbred lines. Notably, Korean mice exhibit a spatially restricted, relatively short and slender, but pointed snout structure [27]. A comprehensive understanding of the evolutionary history of Korean house mice necessitates analyzing both genetic and phenotypic traits across a large sample size.

## Acknowledgments

This research was supported by Basic Science Research Program through the National Research Foundation of Korea (NRF) funded by the Ministry of Education (grant no. 2020R1I1A207315613). This article is dedicated in memory of the late Mr. Jae-Cheol Lee, whose unwavering support was instrumental to this project.

## Supporting information

**S1 Fig. Comparison of craniometric variables between wild house mice from the Middle East** [21] **and our Korean inbred mice.** All craniometric variables depicted in the box plot are represented using a logarithmic scale. Box plot shows mean, range, and minimum and maximum values in the figure. A, DOM; B, CAS (Previously assigned to *M. m. bactrianus* is genetically indistinguishable from CAS [20]); C, MUS; D, *M. m. isatissus*; E, our inbred mice from Goesan-gun.

**S2 Fig. Comparison of craniometric variables of among several inbred mice including our Korean inbred mice.** Box plot shows mean, range, and minimum and maximum values in the figure. The data of A ∼ F are from [22]. A, C57BL/6J; B, BALB/Ca; C, C3H/HeJ; D, CBA/JNCrj; E, ICR; F, MSM/Ms; G, our inbred mice from Goesan-gun.

**S1 Table. Information on samples, GenBank accession numbers for mitochondrial *cytb* sequences.** ^+^ This study. MUS, *M. m. musculus*; CAS, *M. m. castaneus*; DOM, *M. m. domesticus*; MOL, *M. m. molossinus*.

## Notes

### Competing Interest Statement

The authors have declared no competing interest.

